## Supplementary Figures for "Impact of rice *GENERAL REGULATORY FACTOR14h* (*GF14h*) on low-temperature seed germination and its application to breeding"

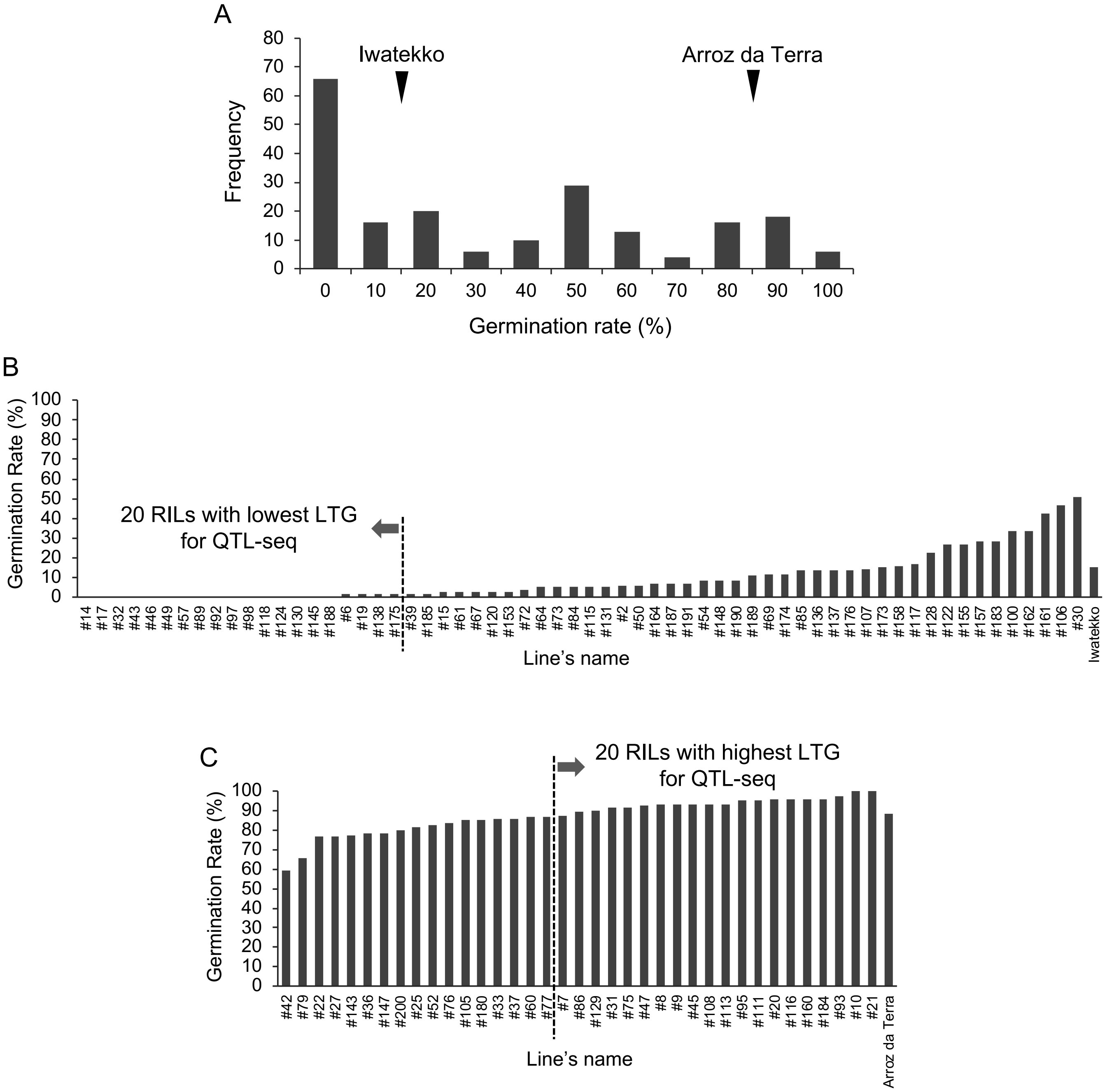


**S1 Fig. Frequency distribution of germination rates in the RIL population and germination rates of selected RILs for QTL-seq analysis.**

**(A)** Frequency distribution of germination rates at $13^{\circ}$ C after eight days from seed imbibition in 200 $F_7$ RILs derived from a cross between Iwatekko and Arroz da Terra [31]. **(B)** Selection of RILs with low cold germination rates. The 62 RILs with the lower germination rates from the first test (A) were tested for the second, and then the 20 RILs with the lowest germination rates were selected as a bulk sample for QTL-seq analysis. The bar graph shows the mean values of the two tests. **(C)** Selection of RILs with high cold germination rates. The 37 RILs with the higher germination rates from the first test (A) were tested for the second, and then the 20 RILs with the highest germination rates were selected for a bulk sample for QTL-seq analysis. The bar graph shows the average values of the two tests.

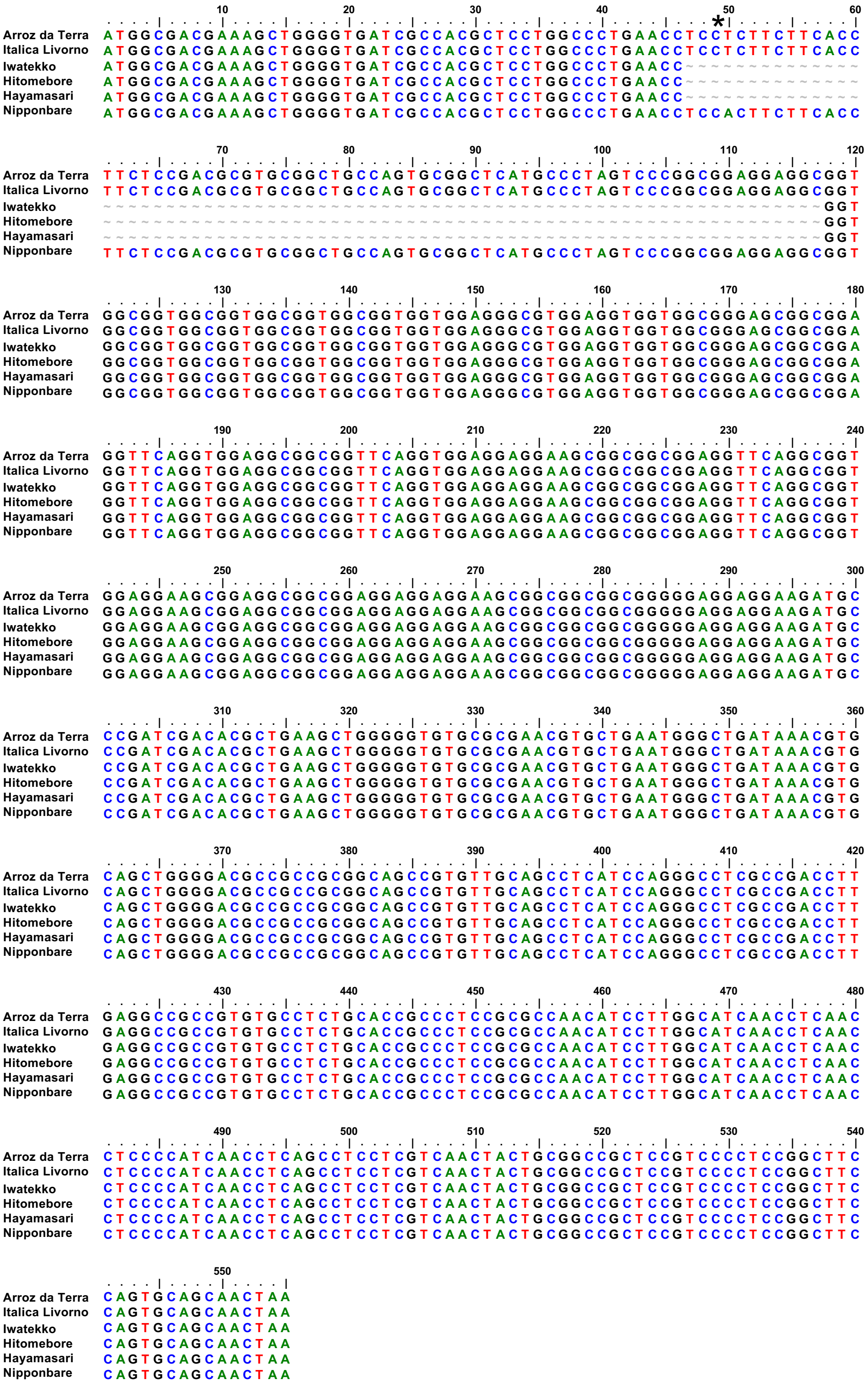


**S2 Fig. Multiple DNA sequence alignment of *qLTG3-1* variants.**
Arroz da Terra and Italica Livorno harbor a functional *qLTG3-1* variant. Nipponbare carries another functional *qLTG3-1* variant due to the nonsynonymous substitution (*). Iwatekko, Hitomebore, and Hayamasari contain a loss-of-function variant for *qLTG3-1* due to a 71-bp deletion.

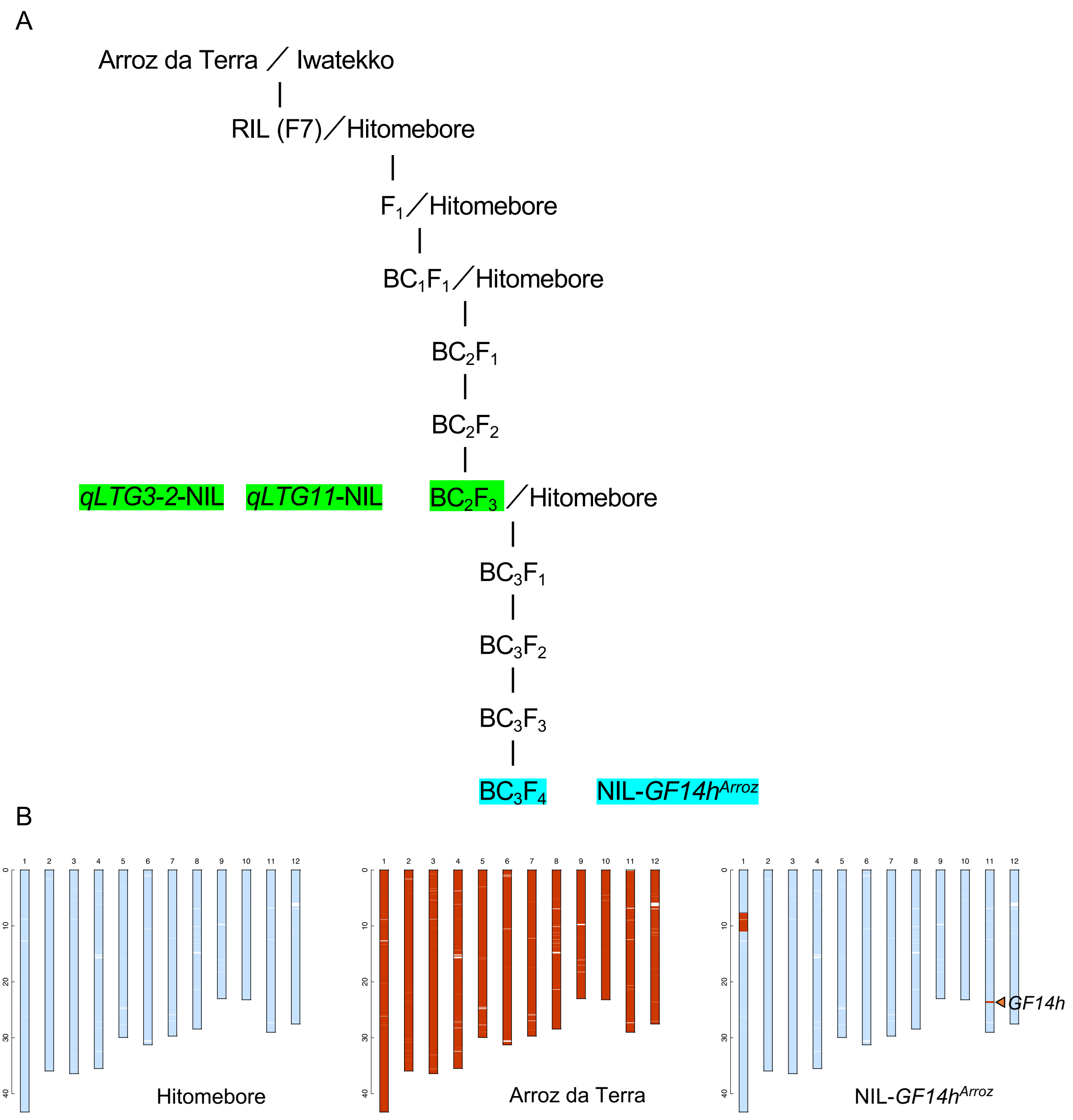


**S3 Fig. Generation of a near-isogenic line with high LTG in the Hitomebore background.**

**(A)** Strategy for the development of *qLTG3-2*-NIL, *qLTG11*-NIL, and NIL-*GF14h*$^{Arroz}$. Molecular markers were used for foreground and background selection. **(B)** Diagram showing the genotype of NIL-*GF14h*$^{Arroz}$. NIL-*GF14h*$^{Arroz}$ contains a 172-kb region on chromosome 11 harboring the Arroz da Terra allele of *GF14h*. Light blue bars indicate genomic fragments from Hitomebore; red bars indicate genomic fragments from Arroz da Terra.

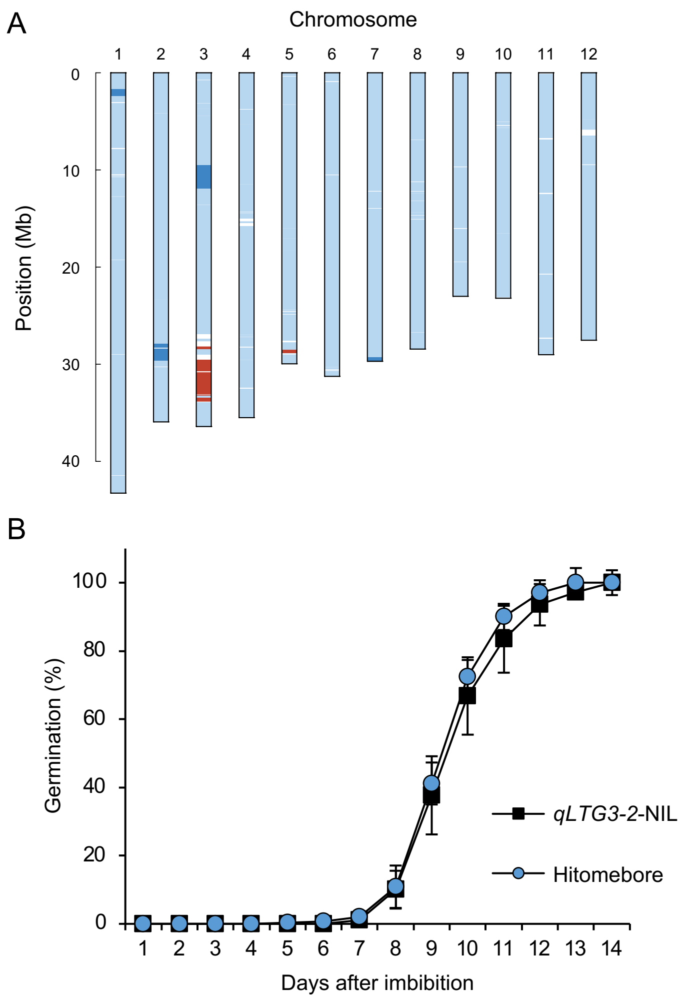


**S4 Fig. Summary of *qLTG3-2*.**

**(A)** Diagram showing the genotype of *qLTG3-2*-NIL containing the Arroz da Terra allele at *qLTG3-2* on chromosome 3. Light blue bars indicate genomic fragments from Hitomebore; red bars indicate genomic fragments from Arroz da Terra; dark blue bars indicate heterozygous regions. **(B)** Germination time courses for seeds of Hitomebore and the *qLTG3-2*-NIL at 15° C. Values are means ± SD of biologically independent samples ($n$ = 10).

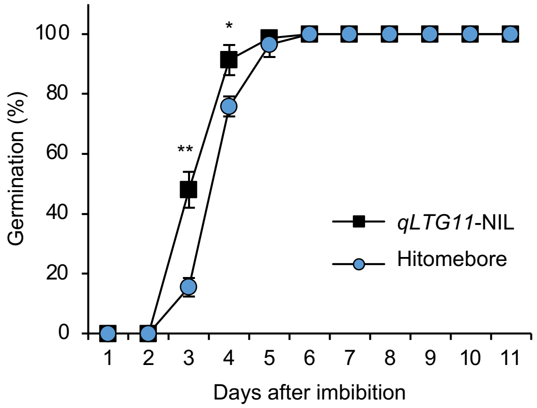


**S5 Fig. Seed germination of *qLTG11*-NIL under optimal temperature conditions.**

Germination time courses of seeds from Hitomebore and *qLTG11*-NIL at 25° C. Values are means ± SD of biologically independent samples ($n$ = 3). Two-tailed t-test was used between *qLTG11*-NIL and Hitomebore for each time point (*$P < 0.05$ and **$P < 0.01$).

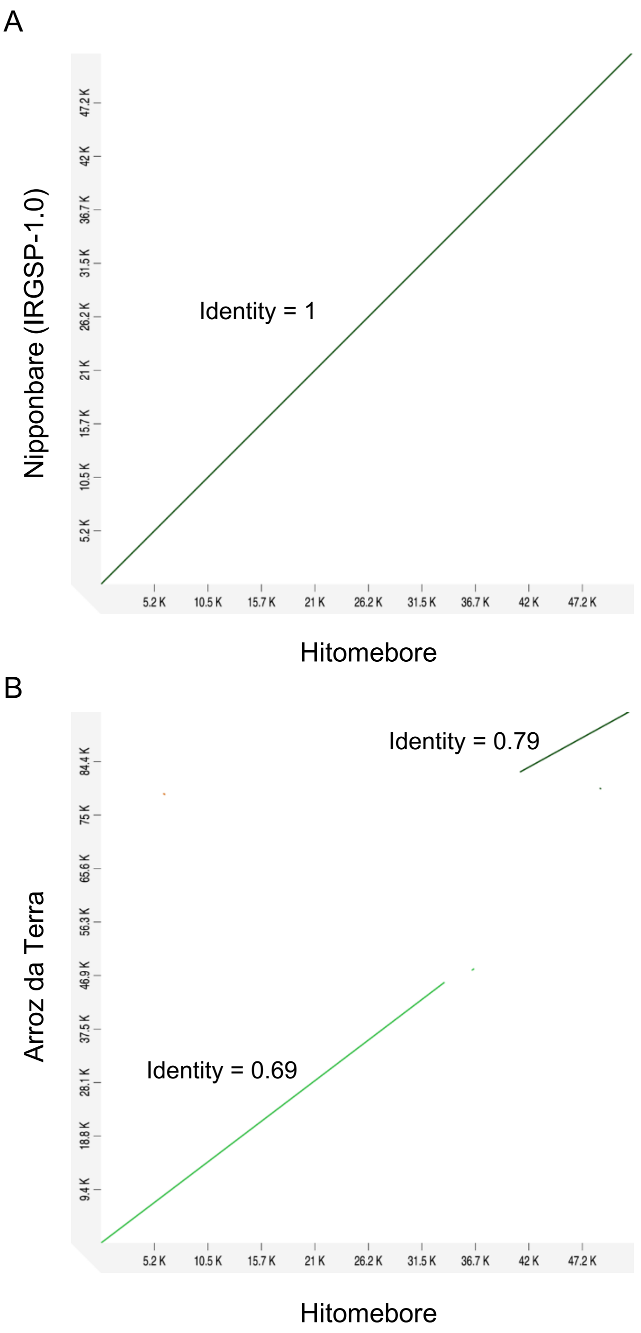


**S6 Fig. Comparison of the *qLTG11* genomic region in Hitomebore, Arroz da Terra, and Nipponbare.**

Dot blot analyses of the genomic sequence in the *qLTG11* candidate region between **(A)** Hitomebore and Nipponbare and **(B)** Hitomebore and Arroz da Terra, using D-GENIES [62]. Based on the Nipponbare genome (IRGSP-1.0), the genomic region containing the causative gene is located at 23.512–23.564 Mb (approximately 52 kb) on chromosome 11. The genome sequence of Hitomebore is identical to that of Nipponbare. The candidate region corresponds to a fragment of approximately 94 kb in the Arroz da Terra genome.

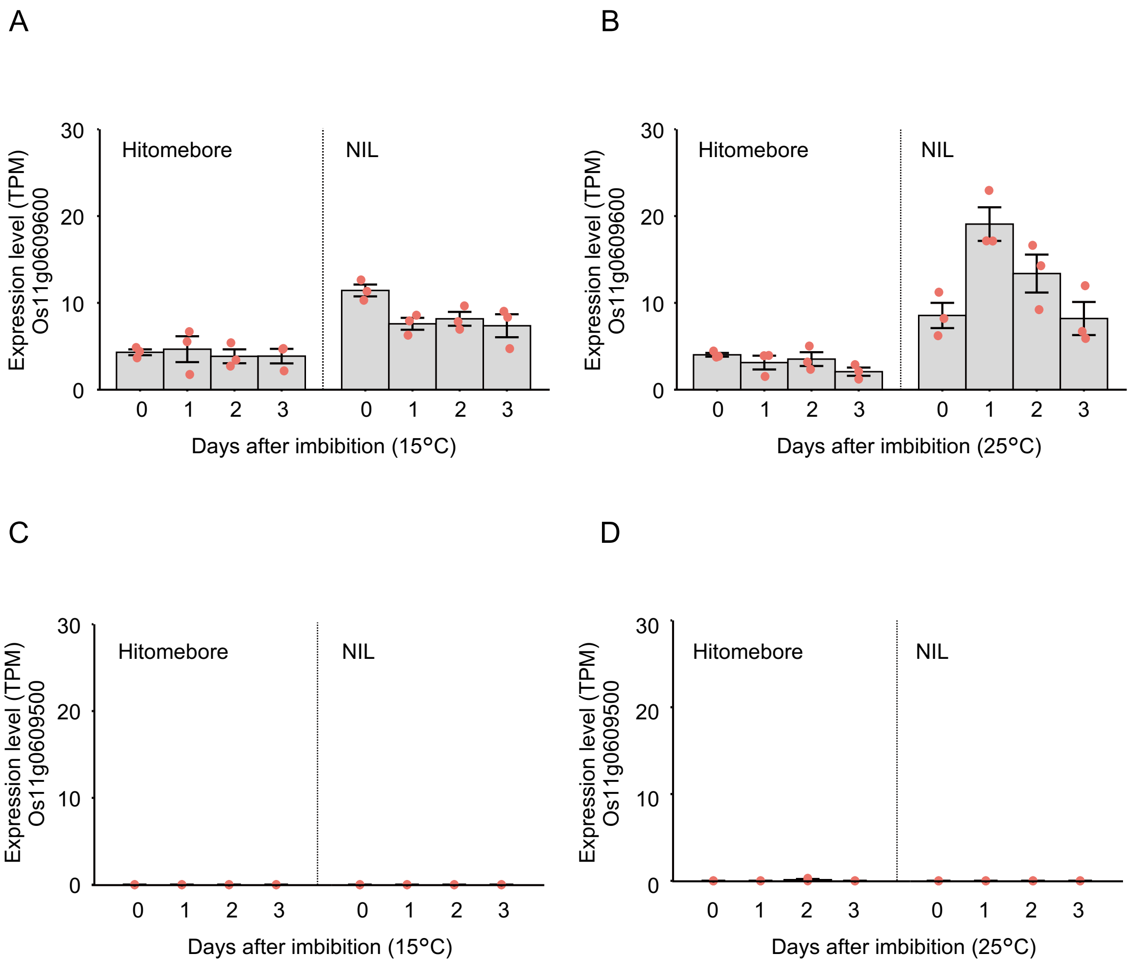


**S7 Fig. Expression levels of the two annotated genes in the candidate genomic region of *qLTG11* based on RNA-seq data.**

Total RNA was extracted from Hitomebore and *qLTG11*-NIL seeds at 0, 1, 2, and 3 days after the onset of seed imbibition under 15 or 25° C temperature conditions, followed by RNA-seq. The sequence reads were mapped to the Nipponbare genome (IRGSP-1.0), and expression data were obtained. **(A–B)** The expression levels of Os11g0609600 (*GF14h*) during seed germination under 15° C (A) and 25° C (B) are shown. Data are presented as means ± SE. n = 3 biologically independent samples. **(C–D)** The expression levels of Os11g0609500 (*Jacalin-like lectin domain containing protein*) during seed germination under 15° C (C) and 25° C (D) are shown. Data are presented as means ± SE. n = 3 biologically independent samples.

```
                       10        20        30        40        50        60
              ....|....|....|....|....|....|....|....|....|....|....|....|
Arroz da Terra ATGAAGGAGAGGGAGAAGGTGGTGCGCCTGGCCAAGCTCGCCGAGCAGGCGGAGAGATAC
Hitomebore     ATGAAGGAGAGGGAGAAGGTGGTGCGACTGGCCAAGCTCGCCGAGCAGGCGGAGAGATAC
Nipponbare     ATGAAGGAGAGGGAGAAGGTGGTGCGACTGGCCAAGCTCGCCGAGCAGGCGGAGAGATAC

                       70        80        90       100       110       120
              ....|....|....|....|....|....|....|....|....|....|....|....|
Arroz da Terra GATGATATGGTGGAATTTATGAAAACACTTGCTAGGATGGATGTGGATATGAGTGCTGAA
Hitomebore     GACGATATGGTGGAATTTATGAAGACACTTGCTAGGATGGATGTGGATATGAGTGCTGAA
Nipponbare     GACGATATGGTGGAATTTATGAAGACACTTGCTAGGATGGATGTGGATATGAGTGCTGAA

                      130       140       150       160       170       180
              ....|....|....|....|....|....|....|....|....|....|....|....|
Arroz da Terra GAGAGGCTTTTATTTTCAGTTGGTTTTAAGAAAACTATTGGTGCCAGGAGAGCATCATGG
Hitomebore     GAGAGGCTTTTATTTTCAGTTGGTTTTAAGAAAACTATTGGTGCCAGGAGAGCATCATGG
Nipponbare     GAGAGGCTTTTATTTTCAGTTGGTTTTAAGAAAACTATTGGTGCCAGGAGAGCATCATGG

                      190       200       210       220       230       240
              ....|....|....|....|....|....|....|....|....|....|....|....|
Arroz da Terra AGAATCCTTGAATCACTCGAGCAGAAGGTGACAGCTGGTGAGCAACCAGGTGTAACAATA
Hitomebore     AGAATCCTTGAATCACTTGAGCAGAAGGTGACAGCTGGTGACCAACCAGGTGTAACAATA
Nipponbare     AGAATCCTTGAATCACTTGAGCAGAAGGTGACAGCTGGTGACCAACCAGGTGTAACAATA

                      250       260       270       280       290       300
              ....|....|....|....|....|....|....|....|....|....|....|....|
Arroz da Terra AATGGCTACAAAAAGAAAGTAGAGGATGAACTAAGGGCAGTTTGCAATGAAGTATTATCA
Hitomebore     AATGGCTACAAAAAGAAAGTAGAGGATGAACTAAGGGCAGTTTGCAATGAAGTATTATCA
Nipponbare     AATGGCTACAAAAAGAAAGTAGAGGATGAACTAAGGGCAGTTTGCAATGAAGTATTATCA

                      310       320       330       340       350       360
              ....|....|....|....|....|....|....|....|....|....|....|....|
Arroz da Terra ATTATTGCTATTCATTGCCTTCCATTGGCCAATTCGGGTGAAAATGTCGTGTTCTTTTAT
Hitomebore     ATTATTGCTATTCATTGCCTTCCATTGGCCAATTCGGGTGAAAATGTCGTGTTCTTTTAT
Nipponbare     ATTATTGCTATTCATTGCCTTCCATTGGCCAATTCGGGTGAAAATGTCGTGTTCTTTTAT

                      370       380       390       400       410       420
              ....|....|....|....|....|....|....|....|....|....|....|....|
Arroz da Terra AAGATGAAGGGTGATTACTACCGTTACTTGGCTGAATTTAGCACTGGAACTGAAAAGAAG
Hitomebore     AAGATGAAGGGTGACTACTACCGTTACTTGGCTGAATTTAGCACTGGAACTGAAAAGAAG
Nipponbare     AAGATGAAGGGTGACTACTACCGTTACTTGGCTGAATTTAGCACTGGAACTGAAAAGAAG

                      430       440       450       460       470       480
              ....|....|....|....|....|....|....|....|....|....|....|....|
Arroz da Terra GCTGCTACTGATCAGTCCCTTATGGCCTATCAGCACGCCATGGTTGTTTCCTCCAGTGAG
Hitomebore     GCTGCTACTGATCAGTCCCTTATGGCCTATCAGCACGCCATGGTTGTTGCCTCCAGTGAG
Nipponbare     GCTGCTACTGATCAGTCCCTTATGGCCTATCAGCACGCCATGGTTGTTGCCTCCAGTGAG

                      490       500       510       520       530       540
              ....|....|....|....|....|....|....|....|....|....|....|....|
Arroz da Terra CTTTCTCCTGTCCATCAATTCAGGCTTGGCCTTGCGCTCAACTTTTCAGTCTTCTTTTAT
Hitomebore     CTTTCTCCTGCCCATCAATTCAGGCTTGGCCTTGCGCTCAACTTTTCAGTCTTCTT----
Nipponbare     CTTTCTCCTGCCCATCAATTCAGGCTTGGCCTTGCGCTCAACTTTTCAGTCTTCTT----

                      550       560       570       580       590       600
              ....|....|....|....|....|....|....|....|....|....|....|....|
Arroz da Terra GAGATAATGAACTCTCCTGAGAGAGCTAGCCAAGTGGCAAAACAAGCACTTGATGAGGCT
Hitomebore     GAGATAA-----------------------------------------------------
Nipponbare     GAGATAA-----------------------------------------------------

                      610       620       630       640       650       660
              ....|....|....|....|....|....|....|....|....|....|....|....|
Arroz da Terra ACTGCTGAGATTAATTCTGCTGGTGTGGAGGGTTACAAGGACAGCATGCTTATGATGCAG
Hitomebore     ------------------------------------------------------------
Nipponbare     ------------------------------------------------------------

                      670       680       690       700       710       720
              ....|....|....|....|....|....|....|....|....|....|....|....|
Arroz da Terra CTCCTGAAGGAGAACCTGGCATTGTGGACATCAGAACTAACTGGAGGTGAAACATCTAAG
Hitomebore     ------------------------------------------------------------
Nipponbare     ------------------------------------------------------------

                      730       740
              ....|....|....|....|...
Arroz da Terra GACGATGACGTTGTTATGGAGGGCTAA
Hitomebore     ---------------------------
Nipponbare     ---------------------------
```

**S8 Fig. Multiple DNA sequence alignment of *GF14h* variants.**

Arroz da Terra carries a functional *GF14h* variant. Hitomebore and Nipponbare harbors a loss-of-function variant of *GF14h* due to a 4-bp deletion (black line).

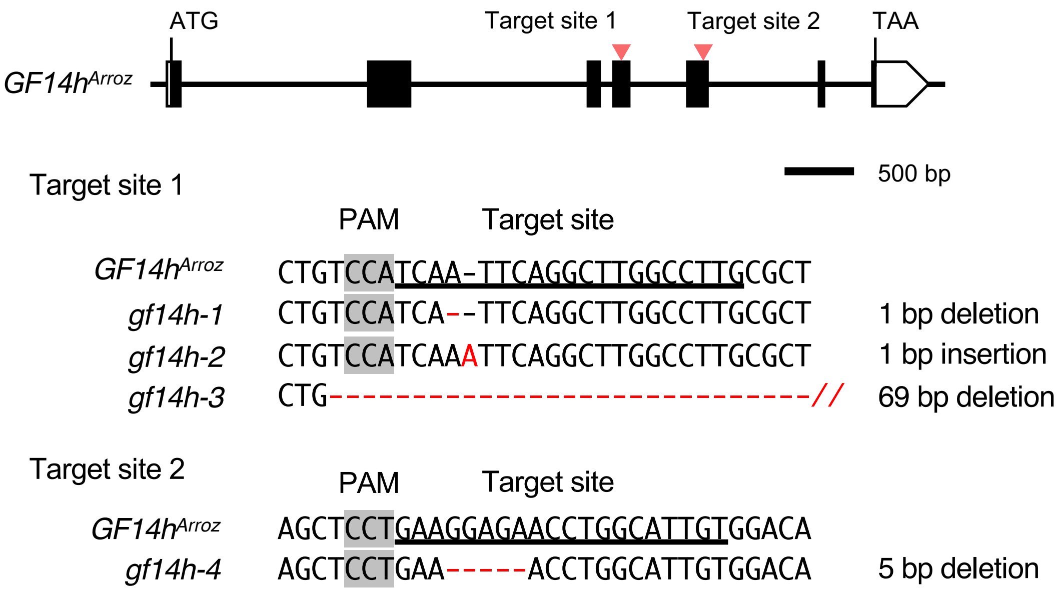


**S9 Fig. CRISPR/Cas9-mediated genome editing of *GF14h*.**

Top, diagram showing the *GF14h* locus, with the locations of the two sgRNA target sites marked by inverted red triangles. Bottom, sequencing results of putative *gf14h* mutants. The sgRNA target sites are underlined, and the PAMs are highlighted. The mutation sites in *GF14h*$^{Arroz}$ for the four mutants (*gf14h-1*, *gf14h-2*, *gf14h-3,* and *gf14h-4*) are indicated.

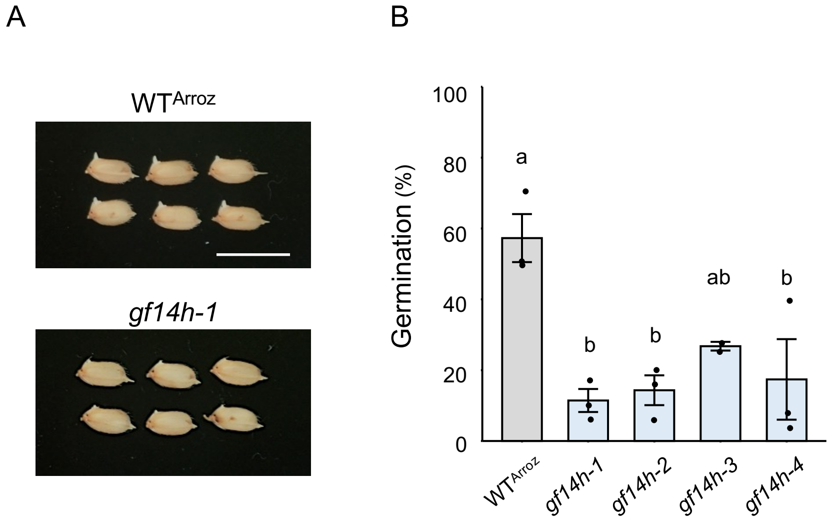


**S10 Fig. Effect of *GF14h* mutation on optimal-temperature germination.**
**(A)** Representative photographs showing seed germination in wild-type harboring Arroz-type *GF14h* ($\mathrm{WT}^{\mathrm{Arroz}}$) and CRISPR/Cas9 knockout lines (*gf14-1*) at 3 days after the onset of seed imbibition. Scale bar, 1 cm. **(B)** Seed germination rate of $\mathrm{WT}^{\mathrm{Arroz}}$ and its CRISPR/Cas9 knockout lines at 2 days of seed imbibition at 25° C. The two target constructs (S9 Fig) were introduced into the *qLTG11*-NIL line. Data are means ± standard error ($\mathrm{WT}^{\mathrm{Arroz}}$, *gf14h-1*, *gf14h-2* and *gf14h-4*, n = 3; *gf14h-3*, n = 2). Different lowercase letters indicate significant differences based on Tukey's HSD test ($P < 0.05$).

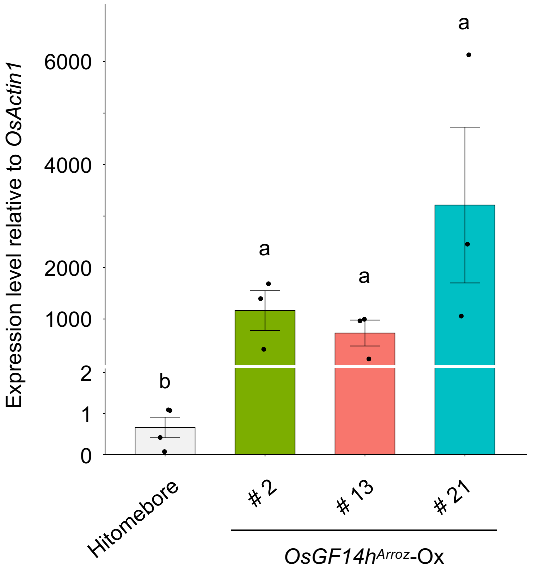


**S11 Fig. Relative *GF14h* expression levels in germinating seeds of *GF14h*$^{Arroz}$ overexpression lines and the parental line.**

The *GF14h*$^{Arroz}$ overexpression construct was introduced into Hitomebore. *OsActin1* (Os03g0718100) was used for normalization. Values are means ± SE (*n* = 3 or 4). Different lowercase letters indicate significant differences based on Tukey's HSD test ($P < 0.001$).

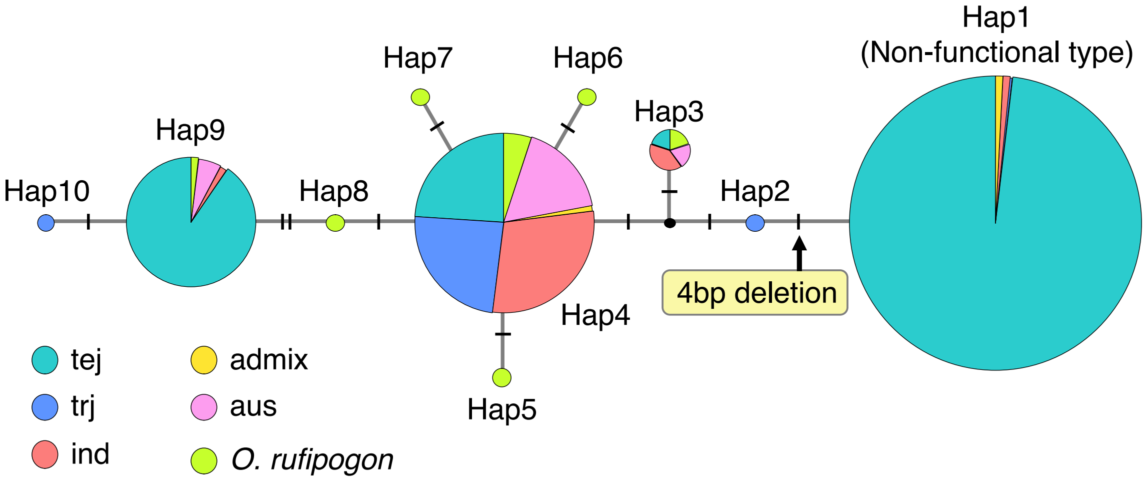


**S12 Fig. Haplotype network of *GF14h*.**

The *GF14h* genomic sequences obtained from 411 *O. sativa* varieties and 11 *O. rufipogon* accessions were used for analysis (S1 Table). The haplotype network was reconstructed by the median joining network algorithm [60] implemented in Popart v1.7 [61]. The haplotype Hap1 evolved from Hap2 by acquiring the 4-bp sequence, resulting in a nonfunctional *GF14h* gene. The Hitomebore cultivar contains Hap1 (nonfunctional), and the Arroz da Terra cultivar contains Hap9 (functional).

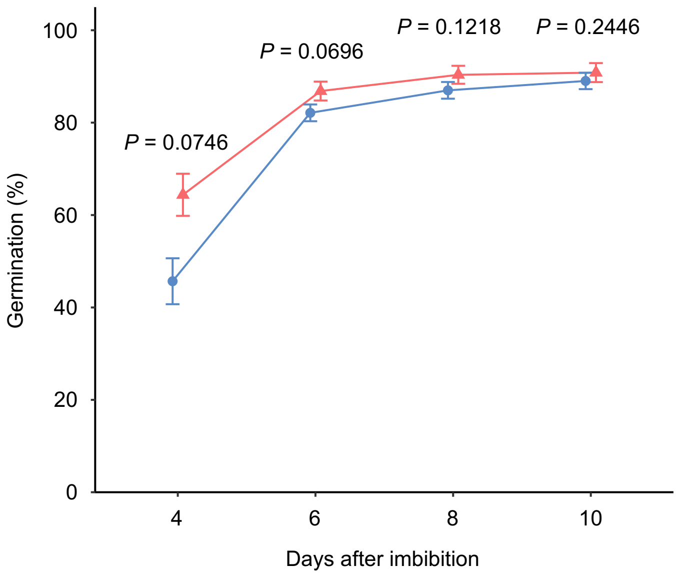


**S13 Fig. Pre-harvest sprouting of NIL-*GF14h*$^{Arroz}$.** Germination time courses of seeds from Hitomebore (blue circles) and the NIL-*GF14h*$^{Arroz}$ (pink triangles) under wet conditions at 28° C. Seeds were harvested from tagged panicles 30 days after heading. Values are means ± SE of biologically independent samples ($n$ = 3). The $P$-values calculated from $t$-tests at each time point are shown in the figure.

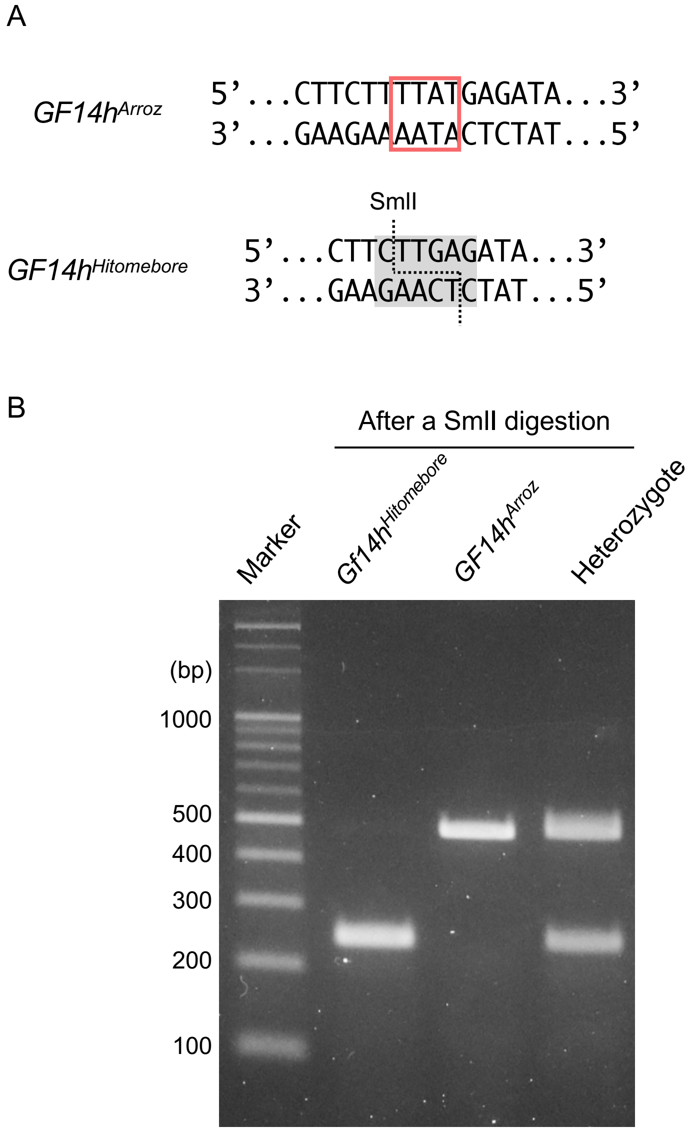


**S14 Fig. Development of a functional marker based on the 4-bp deletion in *GF14h*.**

**(A)** Diagram of the sequence around the 4-bp InDel of *GF14h*. In the Hitomebore (loss-of-function) allele, the 4-bp deletion creates a SmlI restriction site. **(B)** Genotyping of the 4-bp deletion in *GF14h*. A genomic fragment containing the 4-bp InDel of *GF14h* was amplified by PCR and digested with SmlI. The products were separated on a 3% (w/v) agarose gel and stained with Midori Green. The PCR product from *GF14h*$^{Arroz}$ (approximately 500 bp) was not cleaved, whereas the PCR product from *GF14h*$^{Hitomebore}$ was cleaved, producing two fragments of approximately 250 bp each. Both bands were detected in heterozygous plants.
